# Adaptive compensation of arcuate fasciculus lateralization in developmental dyslexia

**DOI:** 10.1101/2022.09.16.508343

**Authors:** Jingjing Zhao, Yueye Zhao, Zujun Song, Michel Thiebaut de Schotten, Irene Altarelli, Franck Ramus

**Author notes:** **Correspondence** Jingjing Zhao, School of Psychology, Shaanxi Normal University, 199 South Chang’an Road, Xi’an, China, 710062, Email address; Franck Ramus, Ecole normale supérieure, Paris.

## Abstract

Previous studies have reported anomalies in the arcuate fasciculus (AF) lateralization in developmental dyslexia (DD). Still, the relationship between AF lateralization and literacy skills in DD remains largely unknown. The purpose of our study is to investigate the relationship between the lateralization of the AF anterior segment (AFAS), AF long segment (AFLS), and AF posterior segment (AFPS) in connection to literacy skills in DD. The participants included 26 children with dyslexia and 31 age-matched children in the control group. High angular diffusion imaging, combined with spherical deconvolution tractography, was used to reconstruct the AF. Connectivity measures of hindrance-modulated oriented anisotropy (HMOA) were computed for each of the three segments of the AF: anterior segment (AFAS), long segment (AFLS), and posterior segment (AFPS). The lateralization index (LI) of each AF segment was calculated by (right HMOA - left HMOA) / (right HMOA + left HMOA). Results showed that the LIs of AFAS and AFLS were positively correlated with reading accuracy in children with dyslexia. Specifically, the LI of AFAS was positively correlated with text and nonword reading accuracy, while the LI of AFLS accounted for word reading accuracy. The results suggest adaptive compensation of arcuate fasciculus lateralization in developmental dyslexia and functional dissociation of the anterior segment and long segment in the compensation.

## Introduction

Developmental dyslexia (DD) is known as a developmental learning disorder with impairment in reading, not caused by a disorder of intellectual development, sensory impairment (vision or hearing), neurological or motor disorder, lack of education availability, lack of proficiency in the language of academic instruction, or psychosocial adversity (World Health Organization, 2018). The prevalence of developmental dyslexia ranges from 1.3% to 17.5% (Di Folco et al., 2020; Shaywitz and Shaywitz, 2005). In the last decade, DD was identified as a disconnection syndrome with defects in the connectivity of the white matter pathways (Gullick & Booth, 2015; Langer, et al., 2017; Saygin, et al., 2013; Su, et al., 2018; Vanderauwera, Wouters, Vandermosten, & Ghesquière, 2017; Vandermosten, et al., 2012).

Arcuate fasciculus (AF) is the most important and well-studied white matter pathway in DD, as it connects two language brain regions, Broca’s and Wernicke’s areas. Several studies reported reduced connectivity indexed by fractional anisotropy (FA) of the left AF in DD compared to the control groups (Christodoulou, et al., 2017; Gullick & Booth, 2015; Langer, et al., 2017; Saygin, et al., 2013; Su, et al., 2018; Vanderauwera, Wouters, Vandermosten, & Ghesquière, 2017; Vandermosten, et al., 2012). In contrast to the deficiency in the left AF, some studies suggest that right AF is related to the reading compensation in DD. In a longitudinal study, Hoeft et al. (2011) reported that FA of the right AF predicted improved reading ability in children with DD over the next 2.5 years. Similarly, studies focusing on children with a family risk of dyslexia have shown that FA of the right AF at pre-school age is correlated with children’s further ability to read (Vanderauwera, Wouters, Vandermosten, & Ghesquière, 2017; Van Der Auwera, Vandermosten, Wouters, Ghesquière & Vanderauwera, 2021; Wang et al., 2017; Zuk et al., 2020). Other research also demonstrates a positive correlation between the FA of the right AF and rapid automatic naming (RAN) in children with dyslexia (El-Sady et al., 2020). These results indicate that the right AF provides considerable compensation in individuals with dyslexia or risk of dyslexia.

Taken together, previous studies suggest a deficiency of the left AF and a compensation of right AF in DD. These results indicate a possibility of atypicality of the left lateralization in the AF and a compensatory role of the right lateralization in the AF. Indeed, atypical left lateralization in the AF has been reported in dyslexic adults: DD showed less left lateralization in AF than the control groups (Vandermosten, Poelmans, Sunaert, Ghesquière, & Wouters, 2013), although we failed to replicate this atypical left lateralization in the AF in dyslexic children (Zhao, Thiebaut de Schotten, Altarelli, Dubois, & Ramus, 2016). In terms of the compensatory role of AF, most previous studies focused on the compensatory role of the right AF in DD. It is unclear whether the right lateralization of the AF can compensate literacy skills in DD. We previously observed that right lateralization of inferior frontal-occipital fasciculus (IFOF) and local network centered at the right fusiform gyrus (FG) could provide maladaptive compensation for literacy skills in dyslexic children (Zhao et al., 2016; Liu, Thiebaut de Schotten, Altarelli, Ramus, & Zhao, 2021). It should be noted that in Zhao et al. (2016) we observed group differences between control and dyslexic children in the right lateralization of IFOF, while in Liu et al. (2021) we found no difference in the local network connectivity of the right FG. However, both right lateralization of IFOF and right FG connectivity were negatively correlated with reading ability of dyslexic children. In particular, right lateralization of IFOF is more associated with word reading ability, while right FG is more associated with pseudoword reading ability. Based on these observations, the present study is intended to test whether compensation exists for the right lateralization of the AF. Specifically, we are intended to test whether right lateralization of the AF is correlated with reading skills, although no significant differences of lateralization in AF were observed between control and dyslexic children.

On the other hand, although compensation of the AF in DD has been reported, not enough research has been done on the compensatory roles different segments of the AF play for different literacy skills in DD. It is well-accepted that the AF can be divided into three distinct segments (AFLS: long frontotemporal, AFAS: anterior frontoparietal, and AFPS: posterior temporoparietal segments) (Catani, Jones, & ffytche, 2005). The long segment, AFLS, connects the language areas in the frontal and temporal lobes. The anterior segment, AFAS, connects the language areas in the frontal and parietal lobes. The posterior segment, AFPS, connects the language areas in the temporal and parietal lobes. Thus, another goal of the present study is to establish whether AFAS, AFLS, and AFPS play different compensatory roles for literacy skills in DD.

In sum, the aim of the present study is twofold. First, whether there is compensation in the lateralization of the AF and its potential adaptive or maladaptive nature. In the case of a positive correlation between the LI of the AF and reading ability, the hypothesis is that the compensation of the AF LI would be adaptive. In the case of a negative correlation between the AF LI and reading ability, the compensation of AF LI would be maladaptive. Second, whether the lateralization of the three segments of the AF has the same compensation mechanism. Specifically, this study explores whether different segments of the AF compensate for the same literacy skills or play different roles in the compensation for different literacy skills (e.g., word reading, pseudoword reading, and text reading).

## Materials and Methods

### Research transparency and Data availability

This study’s experimental procedures and analyses were not registered prior to the research being conducted. The sample size, all the inclusion/exclusion criteria, data exclusions, manipulations, and measures in the study were reported. Inclusion/exclusion criteria were established before the data analysis. Further details can be found in our previous published paper, where we used the same dataset (Zhao, Schotten, Altarelli, Dubois, & Ramus, 2016).

The conditions of our ethics approval do not permit public archiving of anonymized study data. Readers seeking access to the data should contact the lead authors, Franck Ramus and Jingjing Zhao. Access will be granted to named individuals in accordance with ethical procedures governing the reuse of sensitive data. Requestors must provide a research proposal following the guidelines at the Open Science Framework (OSF) approved by Franck Ramus and preregister the study at OSF (https://osf.io) to obtain the data.

### Participants

The study included 26 children with dyslexia, and 31 controls matched for age, gender, handedness, and nonverbal IQ. The age range of the children was 109 - 169 months. All the children’s native language was French. Children with dyslexia were referred from a clinic for reading and language disabilities. They were required to show more than an 18-month delay on reading age based on the accuracy and speed of the Alouette test (Lefavrais, 1967), while the control children had to be no more than 12 months behind. All the subjects had normal hearing and vision abilities no history of brain injury, mental illness, or cognitive impairment. The study was approved by the ethics committee of Bicetre Hospital, and informed consent was obtained from all children and their parents. Analyses of group differences in white matter pathways and white matter network connectivity in the same sample were previously published; no difference in head motion parameters was found between dyslexia and controls (Liu et al., 2021; Lou et al., 2019; Zhao et al., 2016).

### Behavioral measures

We determined the subjects’ intelligence and reading ability through a series of behavioral tests. Intelligence abilities were tested through the WISC blocks, matrices, similarities, and comprehension subtests (Wechsler, 2005). The reading ability test was measured by the Alouette test (Lefavrais, 1967), a test of reading accuracy and speed for meaningless text, and the reading fluency test of words and non-words from Odedys (Jacquier-Roux, Valdois, & Zorman, 2005). Spelling skills were measured by a word spelling-to-dictation test (Martinet & Valdois, 1999).

To analyze the relationship between the brain and the behavior, three composite measures were defined by averaging z scores, according to our previous study (Zhao et al., 2016), as follows: reading accuracy (READACC) was from the word, pseudoword, and text reading accuracy; spelling (SPELL) was simply the z-score of the word spelling test. Positive z-scores were adjusted to signify above-average performance. Legal copyright restrictions prevent publicly archiving various assessment tests used in the current study. They can be obtained from the copyright holders in the cited references.

### Image Acquisition and Analysis

3T MRI scanner (Tim Trio, Siemens Medical Systems, Erlangen, Germany) equipped with a whole-body gradient (40 m T/m, 200 T/m/sec) and a 32-channel head coil were used for all children’s MRI exams at the Neurospin center, Gif-sur-Yvette, France. T1-weighted structural MRI scans were performed using the MPRAGE sequence (acquisition matrix = 230 × 230 × 224, repetition time (TR) = 2,300ms, echo time (TE) = 3.05ms, flip angle = 9^°^, the field of view (FOV) = 230 mm, voxel size = 0.9 × 0.9 × 0.9 mm^3^)

The spin-echo single-shot EPI sequence was utilized for diffusion MRI scans, with parallel imaging (GRAPPA reduction factor 2), partial Fourier sampling (factor 6/8), and bipolar diffusion gradients to reduce geometric distortions. The brain was imaged in full, with an isotropic spatial resolution of 1.7 mm^3^ (matrix size = 128 × 128, the field of view = 218 mm) and 70 interleaved axial slices. Diffusion gradients were applied along 60 uniformly distributed orientations, with a diffusion weighting of b = 1400 sec/mm^2^ (repetition time = 14,000 msec, echo time = 91 msec). In addition, three photographs were taken without any diffusion gradient (b = 0). Each sequence took about 6 minutes, for a total of 18 minutes of acquisition time.

### DTI analysis

The three sequences’ raw DW data were first concatenated into a single data format, using ExploreDTI (http://www.exploredti.com, see Leemans & Jones, 2009); the images were simultaneously recorded and corrected for subject motion and geometrical distortions. A damped Richardson-Lucy algorithm for spherical deconvolution (SD) was used to estimate multiple orientations in voxels containing different populations of crossing fibers (Dell’Acqua et al., 2010). The algorithm parameters were chosen following Thiebaut de Schotten et al (2011).

For whole-brain tractography, per brain voxel with at least one fiber orientation was chosen as a seed voxel. Euler integration with a phase size of 1 mm was used to propagate streamlines from these voxels and for each fiber’s orientation (as described in Dell’Acqua, Simmons, Williams, & Catani, 2013). When entering a region with crossing white matter tracts, the algorithm adopted the least curvature orientation vector (as described in Schmahmann et al., 2007). When a voxel without fiber orientation was reached, or the curvature between two phases exceeded a threshold of 60, the streamlines were halted. In-house software developed with MATLAB v7.8 was used to perform spherical deconvolution, fiber orientation vector estimations, and tractography (The Mathworks, Natick, MA)

TrackVis (http://www.trackvis.org, see Wedeen et al., 2008) was used to dissect each participant’s tracts in their native space. TrackVis allows tract recognition, visualization in three dimensions, and quantitative analysis on each tract. To extract the tracts of interest, a region-of-interest (ROI) approach was used, with the protocol for defining the ROIs for each fiber’s tract based on previous tractography studies; the three segments of the AF being, AFLS: long frontotemporal, AFPS: posterior temporoparietal, and AFAS: anterior frontoparietal segments (Catani, Jones, & ffytche, 2005).

The ROIs were specified on the MNI152 template, provided with the FMRIB Software Library package (FSL, http://www.fmrib.ox.ac.uk/fsl/) to automate some steps of tract dissection and restrict inter-subject variability related to the operator expertise. The Richardson-Lucy Spherical Deconvolution Algorithm was used to calculate the convergence map (CS maps; Dell’Acqua et al., 2006). Advanced Normalization Tools (ANTs, http://www.picsl.upenn.edu/ANTS/) were used to register each subject’s convergence map to the MNI152 prototype that combined affine and diffeomorphic deformations (Avants et al., 2008; Klein et al., 2009). After that, the ROIs identified on the MNI152 template were inverse deformed to bring them to the native space of each participant.

Individual dissections of the tracts were then visually examined and corrected by two anatomists in each participant’s native brain space (JZ and MTS). For each dissected pathway, hindrance-modulated directed anisotropy (HMOA; Dell’Acqua et al., 2013) was extracted and used as a compact measure of fiber density and connectivity characterizing the diffusion properties along with each tract orientation. The critical variable of interest was HMOA, summed across each entire tract.

### Statistical analysis

Statistical analysis was performed using SPSS software (SPSS26, Chicago, IL). We focused on the functional differentiation of the lateralization of the three segments of the AF (AFLS, AFPS, and AFAS) and calculated the lateralization index [HMOA LI = (right HMOA - left HMOA) / (right HMOA + left HMOA). The pearson’s partial correlation between literacy skills and HMOA LI of each segment of the AF was calculated respectively in children with dyslexia and the control group (e.g., SPELL and READACC), after regressing out age, gender, and parental education. Bonferroni correction was adopted for multiple comparisons. Regression analysis was used to confirm the dissociation between LIs of AFLS, AFAS, and reading accuracy in children with dyslexia. We conducted hierarchical linear regression analyses with age, sex, and parental education as controlled variables in the first step. The LIs of AFLS and AFAS were included in the second step. Reading accuracy was incorporated as a dependent variable. In multiple linear regression, under the control of age, sex, and parent’s education, the relationship between the LIs of AFLS and AFAS relative to the three dimensions of reading accuracy (word reading, pseudoword reading, and text reading) were explored in the dyslexia group, respectively.

## Results

### Demographics and behavioral results

Table 1 shows descriptive statistical information on all participants’ demographics and behavioral performance. There was no difference in age, sex, handedness, and non-verbal IQ between the group with dyslexia and the control groups. However, children with dyslexia performed worse than the control group in all aspects of literacy skills, including word, pseudoword, text reading, and spelling ability.

**Table 1.**
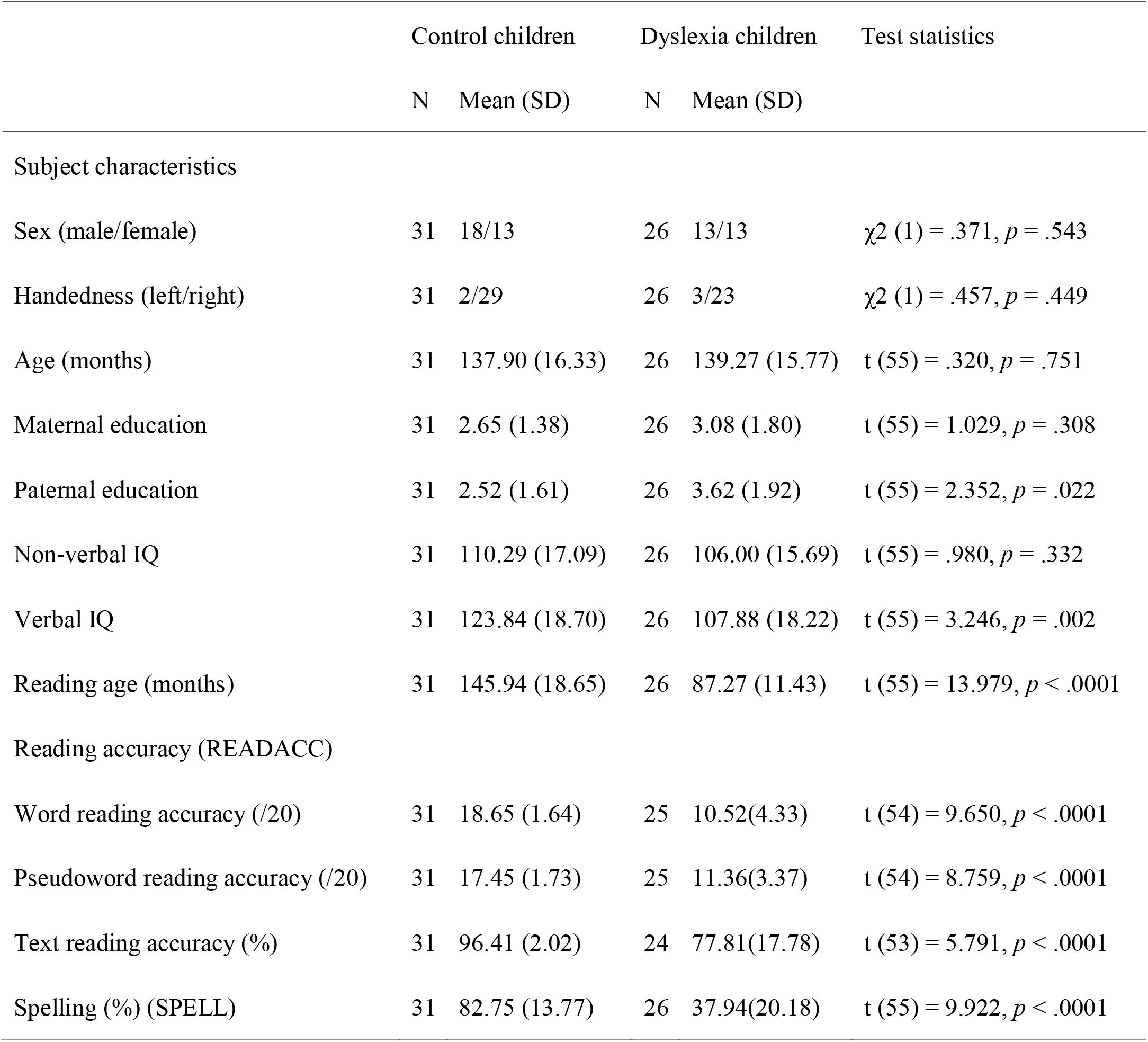
Demographical data and scores of literacy skills.

### Correlation between LI of AF and reading ability

Partial correlation analysis showed a significant correlation between LI of AFAS and reading accuracy in dyslexic children: the more right-lateralized the AFAS, the better reading accuracy in children with dyslexia (*r* = 0.650, *p* = 0.001, see Table 2 and Figure 1). A similar trend was observed in AFLS LI and reading accuracy in dyslexic children (*r* = 0.518, *p* = 0.013), but this result did not survive Bonferroni correction. Linear regression analysis showed that when age, sex, and parents’ education are controlled, the association between AFAS LI and reading accuracy (β= 0.590, *p*= 0.002), as well as the association between AFLS LI and reading accuracy (β=0.373,*p*= 0.020) were significant (see Table 3). This suggests that the functional role of AFAS and AFLS might be independent.

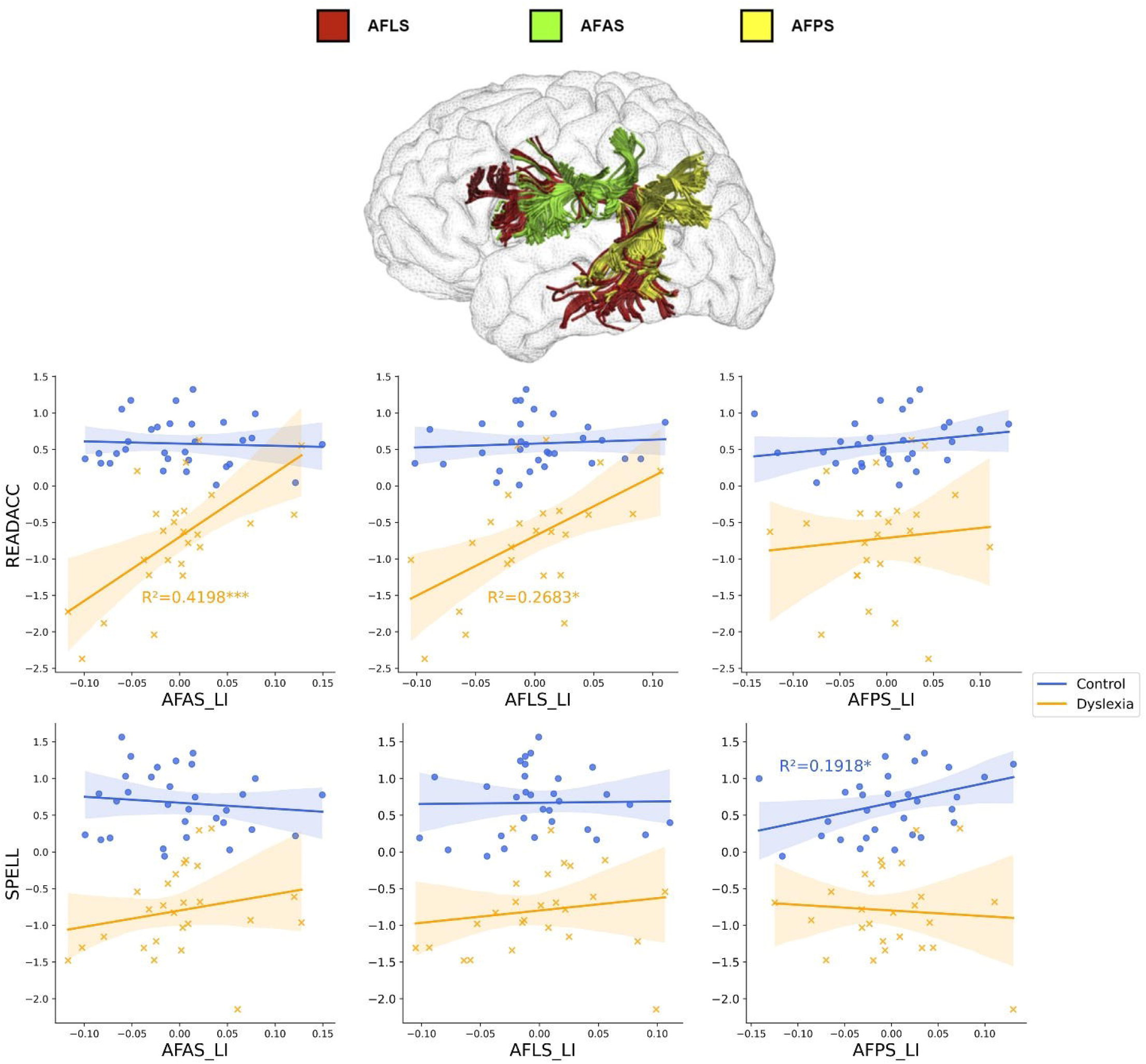

**Table 2.** Partial correlation coefficients (controlled for age, sex, and parental education) between lateralization index (LI) of three segments of arcuate fasciculus (anterior segment: AFAS, long segment: AFLS, and posterior segment: AFPS) and behavioral measures of reading accuracy (READACC), and spelling accuracy (SPELL). ^#^*p* < 0.1, \**p* < 0.05, \*\**p* < 0.01, \*\*\**p* < 0.0042 (surviving Bonferroni correction 0.05/12).

|  |  | AFAS_LI |  | AFLS_LI |  | AFPS_LI |  |
| --- | --- | --- | --- | --- | --- | --- | --- |
|  |  | r | *p* | r | *p* | r | *p* |
| READACC | Dyslexia | **0.650<sup>\*\*\*</sup>** | **0.001** | **0.518<sup>\*</sup>** | **0.013** | 0.038 | 0.867 |
|  | Control | -0.093 | 0.636 | 0.129 | 0.513 | 0.369 | 0.054 |
| SPELL | Dyslexia | 0.377<sup>#</sup> | 0.084 | 0.402<sup>#</sup> | 0.064 | 0.153 | 0.495 |
|  | Control | -0.139 | 0.482 | 0.023 | 0.907 | **0.438<sup>\*</sup>** | **0.020** |

**Table 3.**
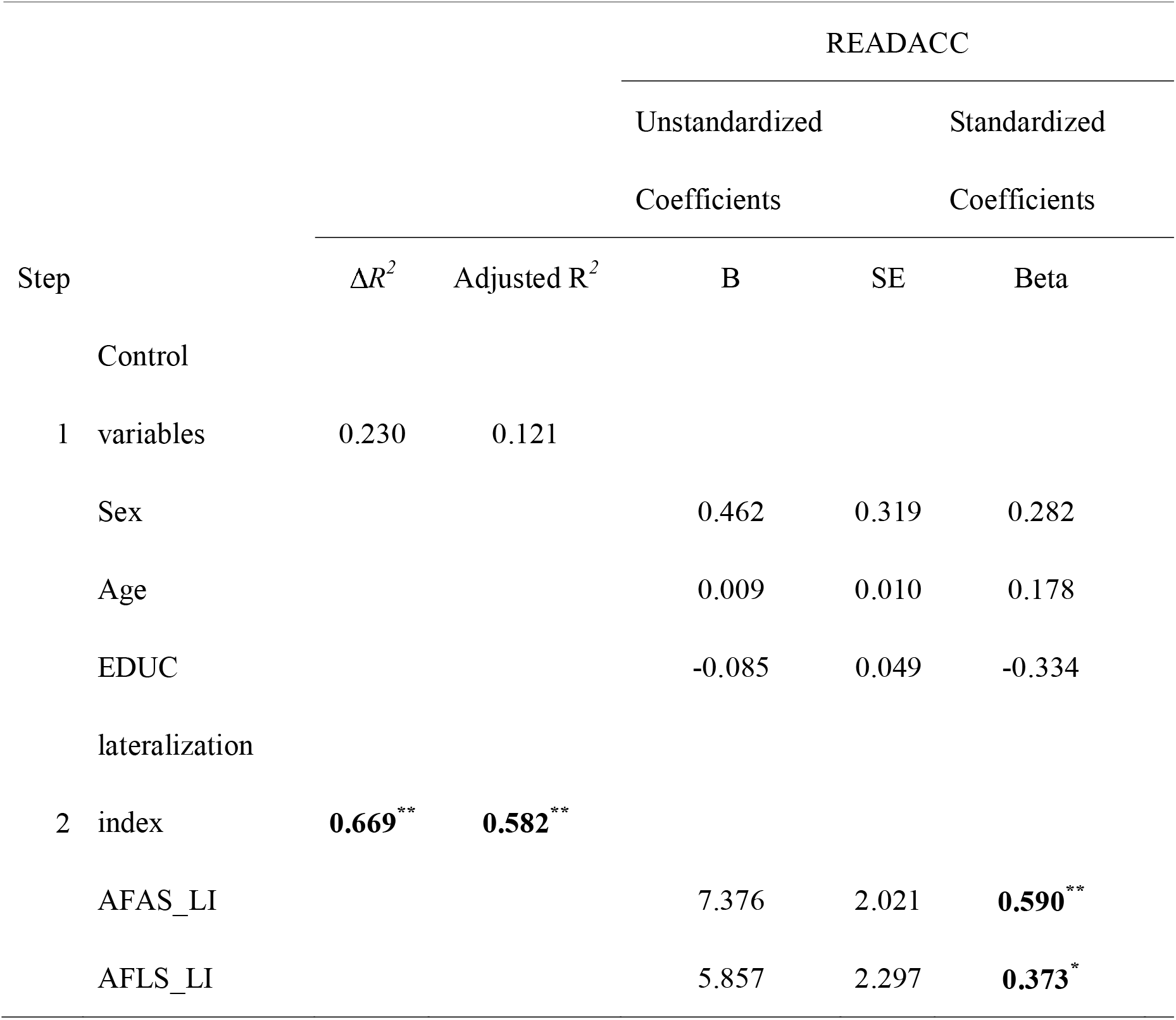
Hierarchical linear regression analysis of reading accuracy (READACC) predicted by the lateralization index (LI) of arcuate fasciculus long segment (AFLS) and arcuate fasciculus anterior segment (AFAS) with age, sex, and parental education controlled in children with dyslexia. \**p* < 0.05, \*\**p* < 0.01.

Three dimensions of reading abilities (word reading, pseudoword reading, and text reading) were further analyzed in reading accuracy in a series of regression analyses (Table 4, 5, and 6). The results of these regressions showed that AFLS is significantly related to word reading accuracy (β= 0.453, *p*= 0.014), but not to pseudoword reading accuracy and text reading accuracy. In contrast, AFAS is significantly related to pseudoword reading accuracy (β= 0.546, *p*= 0.024) and text reading accuracy (β= 0.611, *p*= 0.008), but not to word reading accuracy. Figure 2 shows individual scatterplots for the relationship between the three dimensions of reading accuracy (word reading, pseudoword reading, and text reading) and LIs of AFLS/AFAS. In the dyslexia group, AFLS was related to word reading accuracy (*r* = 0.481, *p* = 0.032), while AFAS was related to pseudoword reading accuracy (*r* = 0.450, *p* = 0.047) and text reading accuracy (*r* = 0.617, *p* = 0.004).

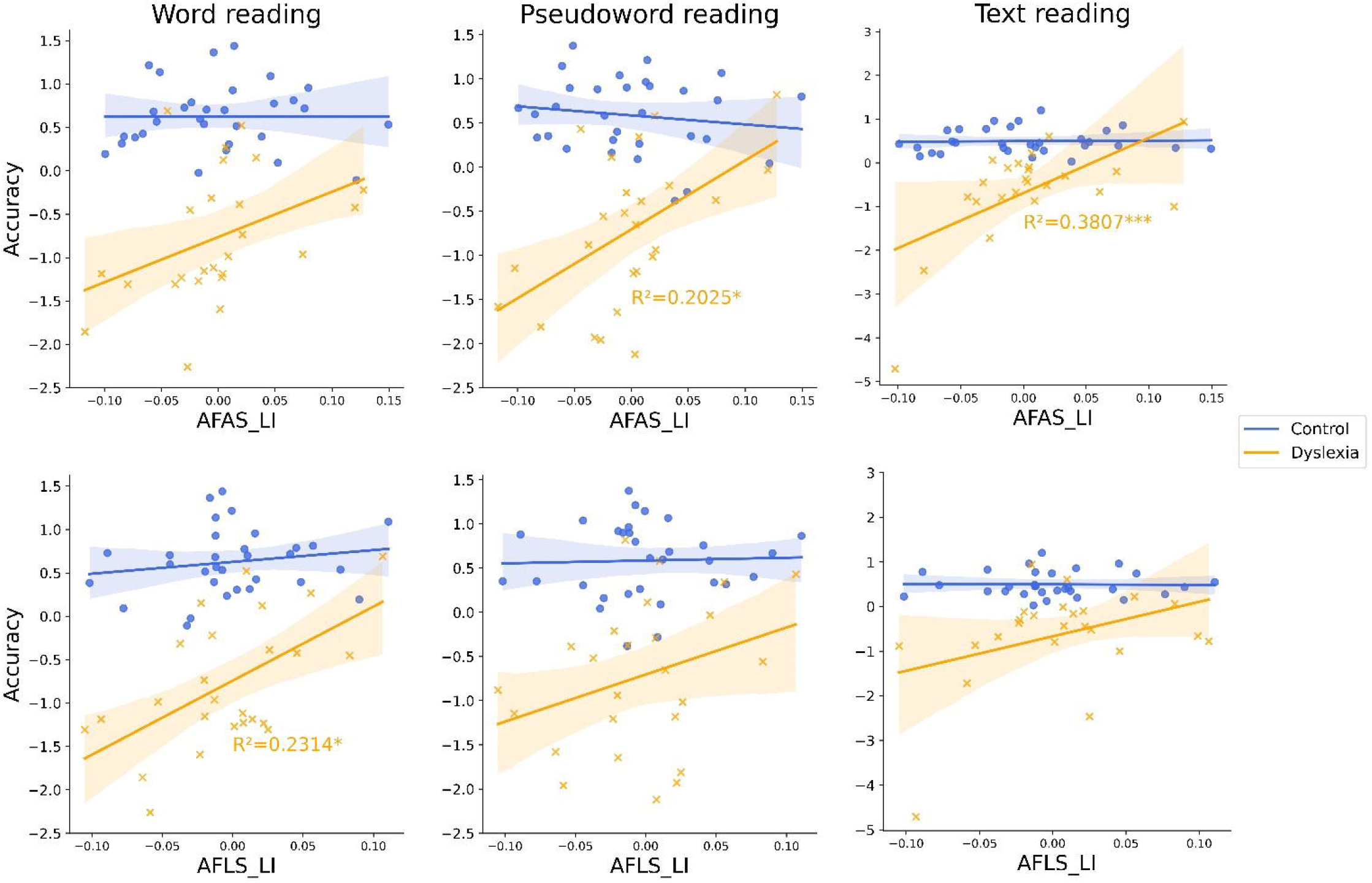

**Table 4.**
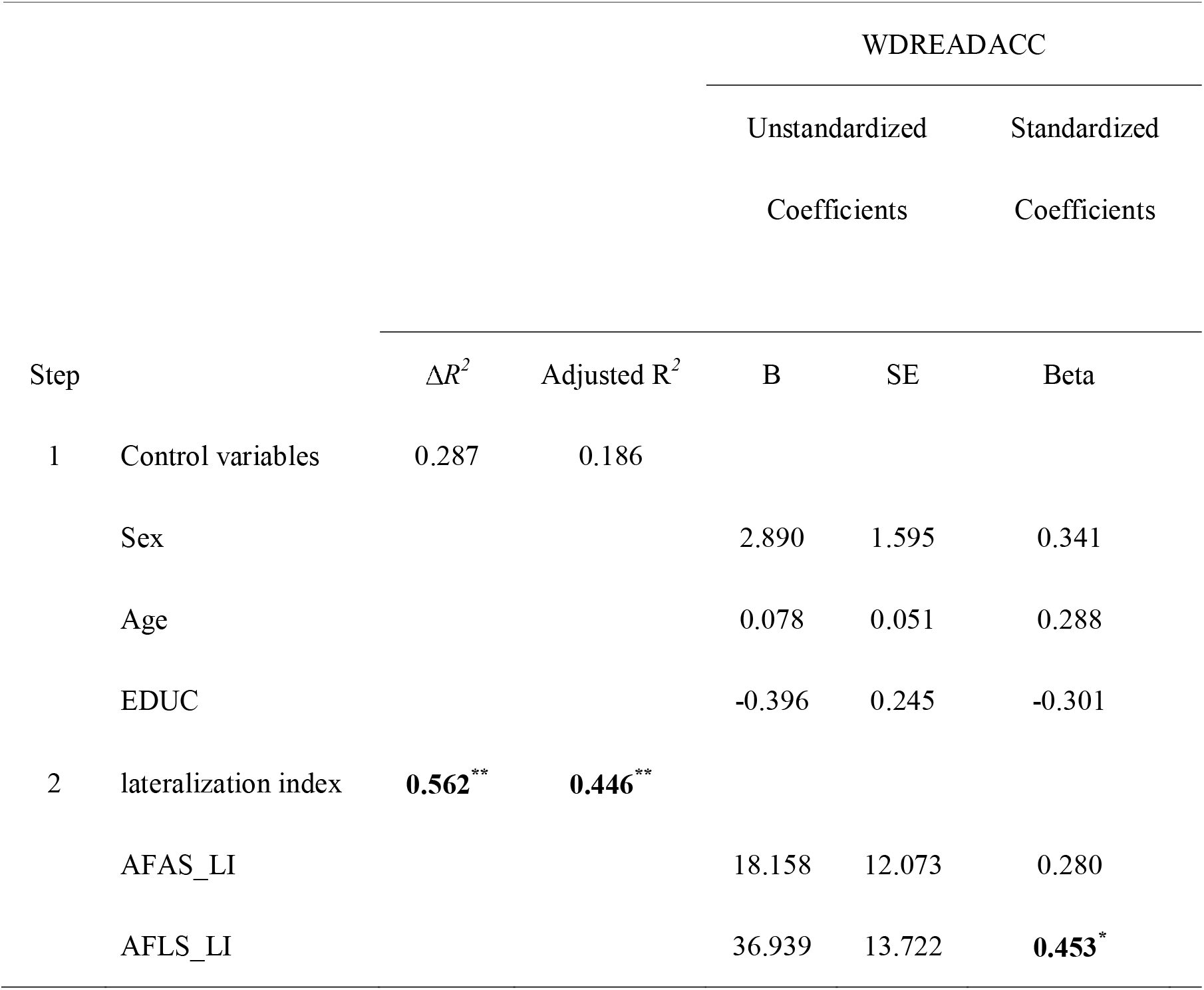
Hierarchical linear regression analysis of word reading accuracy (WDREADACC) predicted by the lateralization index (LI) of arcuate fasciculus long segment (AFLS) and arcuate fasciculus anterior segment (AFAS) with age, sex, and parental education controlled in children with dyslexia. \**p* < 0.05, \*\**p* < 0.01.

**Table 5.**
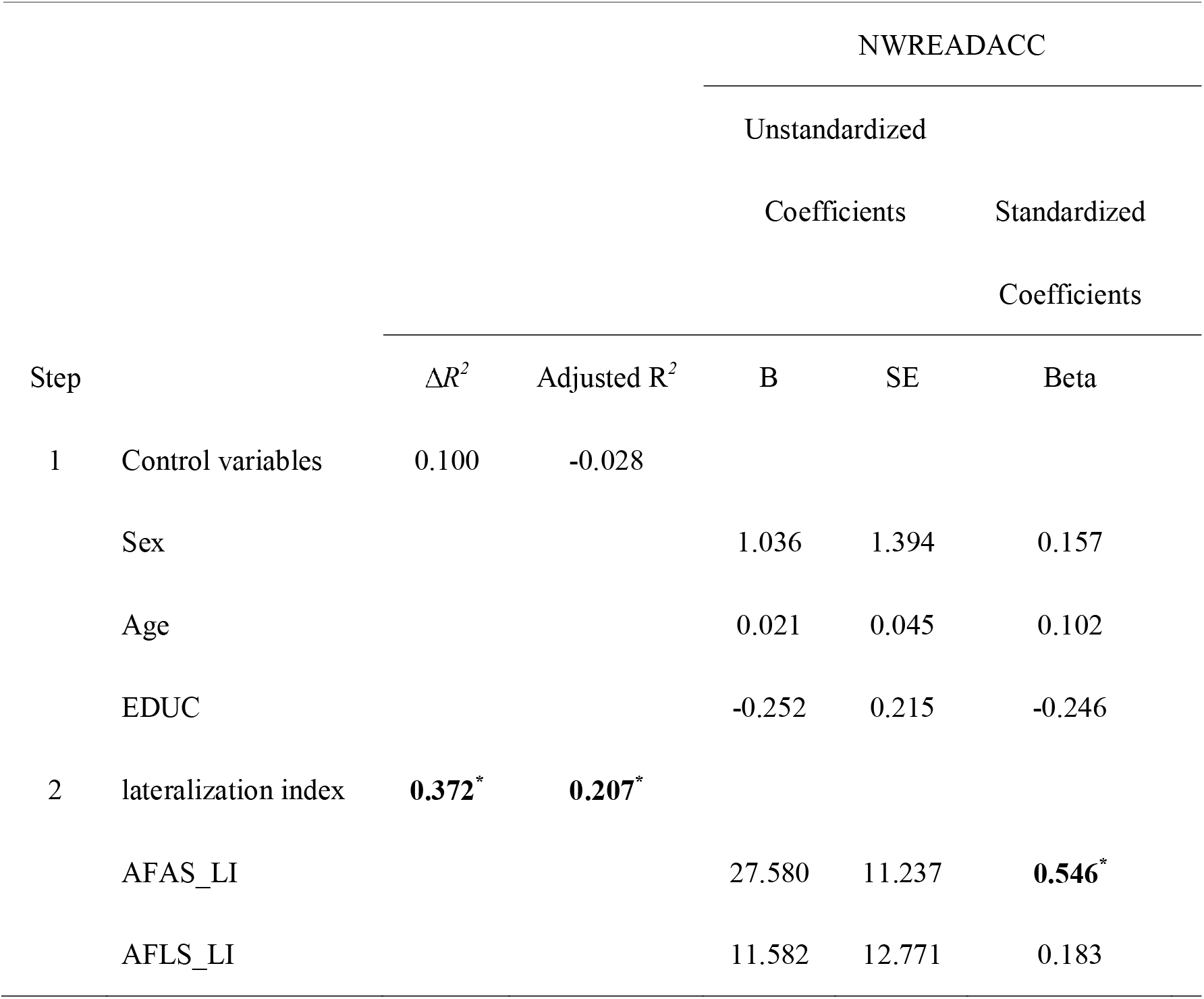
Hierarchical linear regression analysis of pseudoword reading accuracy (NWDREADACC) predicted by the lateralization index (LI) of arcuate fasciculus long segment (AFLS) and arcuate fasciculus anterior segment (AFAS) with age, sex, and parental education controlled in children with dyslexia. \**p* < 0.05.

**Table 6.**
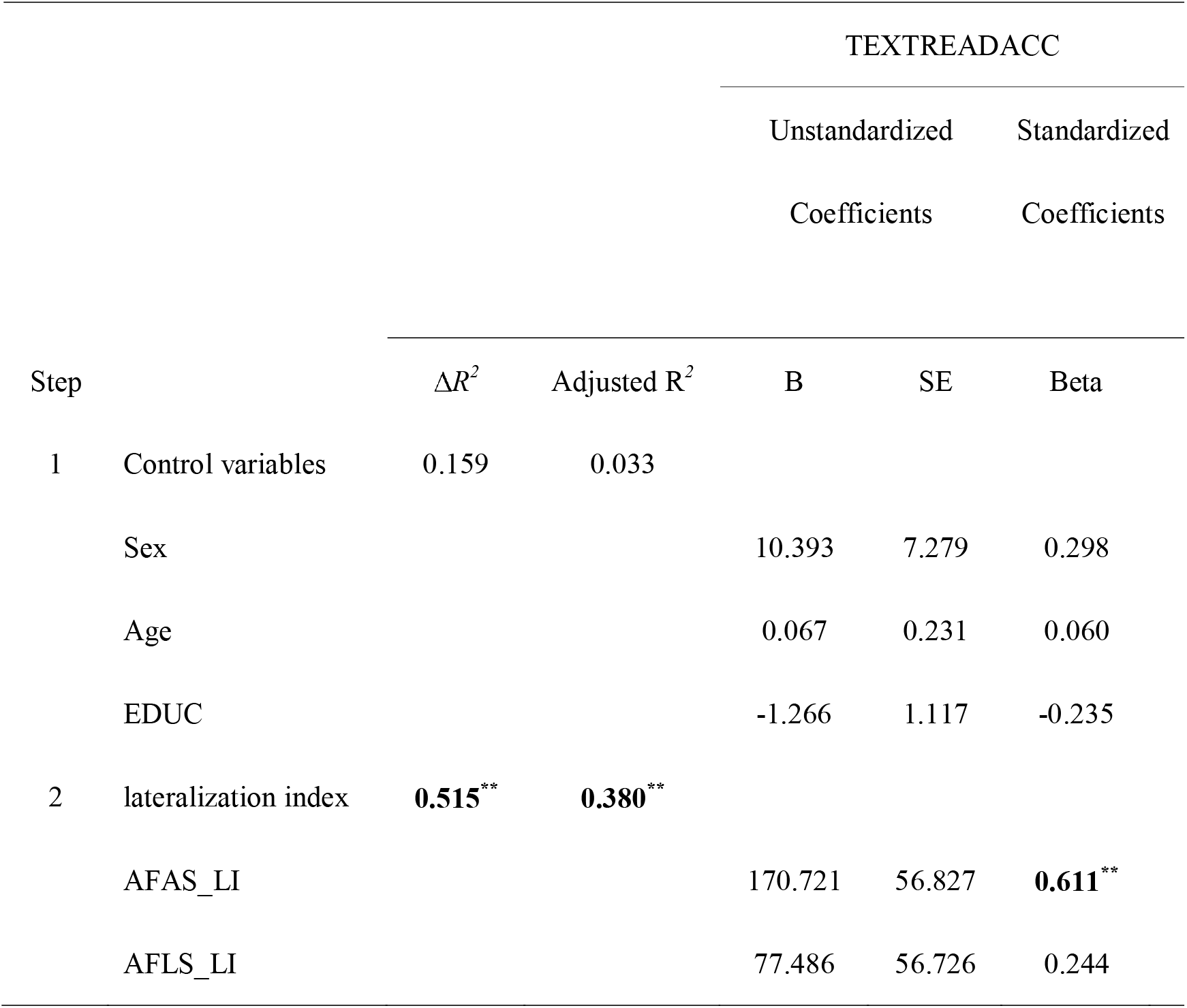
Hierarchical linear regression analysis of text reading accuracy (TEXTREADACC) predicted by the lateralization index (LI) of arcuate fasciculus long segment (AFLS) and arcuate fasciculus anterior segment (AFAS) with age, sex, and parental education controlled in children with dyslexia. \*\**p* < 0.01.

## Discussion

The present study revealed two main facts. First, the results showed a significant positive correlation between the rightward lateralization of the AF and reading accuracy, suggesting a compensatory role of the right AF lateralization. Secondly, we found that the compensatory roles of AFAS and AFLS are functionally independent. The lateralization of AFLS plays a role in word reading accuracy, while the lateralization of AFAS plays a role in pseudoword and text reading accuracy.

Firstly, the results show a positive correlation between the rightward lateralization of the AF and reading skills in dyslexic children. These correlations were only shown between reading skills and LI of AFAS/AFLS, but not between reading skills and the LI of AFPS. These results are consistent with the previous findings that suggest the compensation of right AF in DD or children with risk of DD (Hoeft, et al., 2011; Vander Stappen, Dricot & Van Reybroeck, 2020; Vanderauwera, Wouters, Vandermosten, & Ghesquière, 2017; Wang, et al., 2017). For instance, Hoeft et al (2011) reported that integrity of the right SLF, including AF anterior and long segments, could predict long-term improvement of the reading performance in children with dyslexia. Thus, the stronger the integrity of the right AF SLF, the better the reading level over the next two and a half years. However, Hoeft and colleagues did not further anatomize the right SLF/AF into more refined anterior and long segments of the AF. This paper provides the first account of the independent compensation role of the right lateralization of AFAS and AFLS (Table 3). Our findings, together with the previous results, suggest that the impairment of the left AFAS and AFLS in dyslexia can be repaired or compensated by the right AF through later development or training. It should also be noted that the present study reports the association between rightward AF lateralization and reading skills in DD for the first time. Secondly, it is essential to emphasize the following finding: AFLS and AFAS compensate for different aspects of reading abilities. Generally, the lateralization of AFLS compensates for word reading accuracy, while the lateralization of AFAS compensates for meaningless text reading and nonword reading ability. Previous studies have suggested that AFLS is related to phonological processing, such as phonemic awareness, language fluency, and verbal working memory, while AFAS is related to the phonological storage and articulatory rehearsal module (Duffau, 2008; Vandermosten et al., 2012; Li, et al., 2017; Meyer, Cunitz, Obleser, & Friederici, 2014; Nakajima, Kinoshita, Shinohara, & Nakada, 2020; Papagno, et al., 2017). The present results propose that the right lateralization of AFLS might be related to the compensation of phonological processing of the words.

In contrast, the right lateralization of AFAS might be associated with the compensation of speech articulation of the words. These data provide further evidence for understanding the compensation functions of the AF. They support that different segments of the AF might represent compensatory neural bases related to different aspects of the phonological deficit in dyslexia (Ramus, 2001; Ramus, 2003; Ramus & Szenkovits, 2008).

Lastly, in contrast to our previous findings on maladaptive compensation (Zhao et al., 2016; Liu et al., 2021), this study represents the first demonstration of an adaptive compensation in dyslexia. Previous studies of our group reported negative correlations between the rightward lateralization of inferior frontal-occipital fasciculus (IFOF) and reading accuracy (Zhao et al., 2016), as well as between the right fusiform and reading accuracy (Liu et al., 2021). Similarly, Banfi et al. (2019) reported a negative correlation between the right inferior longitude fasciculus (ILF) and pseudoword reading. These findings point to a maladaptive compensation network in the ventral language pathway. Comparatively, this adaptive compensation pathway, observed in the AF in the present study, was in the dorsal language network. Therefore, these results might suggest that different neural pathways correspond to different compensation modes: adaptive compensation in the dorsal language pathway and maladaptive compensation in the ventral language pathway.

Finally, it should be acknowledged that our study is a cross-sectional correlation study. Thus, no longitudinal follow-up was conducted, unlike in the previous studies (Hoeft, et al., 2011; Zuk, et al., 2021). Therefore, it was impossible to explore the long-term prediction of the AF compensation lateralization for reading ability in DD. In addition, given the limited dyslexia sample in each age group across 9-14 years in our study, the age effect of the AF lateralization compensation was not examined. These are the suggested areas of future research.

In summary, our study revealed an adaptive compensatory role of the AF lateralization in developmental dyslexia. Additionally, a different adaptive compensatory role was found for the long and anterior segments of the AF. The long segment of the AF is compensatory for word reading accuracy, whereas the anterior segment of the AF is compensatory for pseudoword and text reading accuracy.

## Credit Author Statement

Jingjing Zhao conceived of the presented idea. Irene Altarelli and Franck Ramus collected the behavioral, MRI, and diffusion tensor imaging data. Jingjing Zhao, Yueye Zhao and Zujun Song analyzed the data and wrote the main manuscript. Michel Thiebaut de Schotten supervised the analytical methods and revised the manuscript. Franck Ramus verified the analytical methods, supervised the findings, and revised the manuscript.

## Acknowledgments

National Natural Science Foundation of China funded the study (61807023), Humanities and Social Science Fund of Ministry of Education of the People’s Republic of China (17XJC190010), Shaanxi Province Natural Science Foundation (2018JQ8015), and Fundamental Research Funds for the Central Universities (CN) (GK201702011). The study was also funded by Agence Nationale de la Recherche (ANR-06-NEURO-019-01, ANR-11-BSV4-014-01, ANR-17-EURE-0017 and ANR-10-IDEX-0001-02), Commission of the European Communities (LSHM-CT-2005-018696), École des Neurosciences de Paris Île-de-France. MTS received funding from the European Research Council (ERC) under the European Union’s Horizon 2020 research and innovation programme (grant agreement no. 818521). The authors would like to thank all children who participated in the study and their parents, for their time and cooperation. We acknowledge the collaboration of Catherine Billard, Stéphanie Iannuzzi, Nadège Villiermet, Ghislaine Dehaene-Lambertz and all the staff at Neurospin for recruitment and testing.

